# Genome-wide peptidoglycan profiling of *Vibrio cholerae*

**DOI:** 10.1101/2022.08.25.505259

**Authors:** Sara B. Hernandez, Laura Alvarez, Barbara Ritzl-Rinkenberger, Bastian Schiffthaler, Alonso R. Serrano, Felipe Cava

## Abstract

Most bacteria cells are protected by a peptidoglycan cell wall. Defining the chemical structure of the peptidoglycan has been instrumental to characterize cell wall associated proteins and to illuminate the mode of action of cell wall-acting antibiotics. However, a major roadblock for a comprehensive understanding of peptidoglycan homeostasis has been the lack of methods to conduct large-scale, systematic studies. Here we have developed and applied an innovative high throughput peptidoglycan analytical pipeline to analyze the entire non-essential, arrayed mutant library of *Vibrio cholerae*. The unprecedented breadth of these analyses revealed that peptidoglycan homeostasis is preserved by a large percentage of the genome organized in complex networks that functionally link peptidoglycan features with genetic determinants. As an example, we discovered a novel bifunctional penicillin-binding protein in *V. cholerae*. Collectively, genome-wide peptidoglycan profiling provides a fast, easy, and unbiased method for systematic identification of the genetic determinants of peptidoglycan synthesis and remodeling.

## INTRODUCTION

Most bacteria present a strong, yet elastic, cell wall surrounding the cytosolic membrane that maintains cell turgor, determines the bacterial shape and serves as scaffold for the attachment of other cellular components (Dufresne and Paradis-Bleau, 2015). The main component of the bacterial cell wall is the peptidoglycan (PG), a polymer of glycan chains of the alternating *N*-acetyl-glucosamine (NAG) and *N*-acetyl-muramic acid (NAM) disaccharide that are crosslinked through stem peptides. PG composition and structure can vary between bacterial species and also in response to environmental changes (i.e., growth conditions) (Vollmer et al., 2008).

PG is a unique and essential component in bacteria, and hence the enzymes that synthesize and remodel this polymer are commonly used as antibacterial targets. Consequently, extensive research has been performed during decades to provide a more comprehensive understanding of the genetic and environmental factors that govern PG homeostasis (Dorr, 2021; Hernandez and Cava, 2021; Kumar et al., 2022; Porfirio et al., 2019; Rohs and Bernhardt, 2021). Unfortunately, conventional methods for the fine analysis of the chemical structure of the PG are tedious, time-consuming and require a significant culture size as starting material, making them largely incompatible with large-scale, systematic screenings.

Current knowledge about the bacterial PG chemical structure has been mainly obtained by analytical methods proposed by Glauner *et al.* more than 30 years ago (Glauner et al., 1988), which involves: purification of the mature PG sacculus, removal of proteins covalently linked to the PG, digestion with muramidase (i.e., lysozyme) to generate soluble disaccharide peptides (muropeptides) and separation of the muropeptides by liquid chromatography (LC) (Alvarez et al., 2016; Desmarais et al., 2013). These analyses have significantly improved during the last decade thanks to the use of a more advanced ultra-performance liquid chromatography (UPLC) technology replacing high pressure liquid chromatography (HPLC) systems, and the use of in-line LC-mass spectrometry (MS) systems for PG analysis (Alvarez et al., 2016; Bern et al., 2017; Desmarais et al., 2013; Kuhner et al., 2014; Patel et al., 2021). Additionally, the PG isolation protocol has also been simplified and accelerated by replacing the use of the ultra-centrifuge by bench centrifuges, and the reduction or removal of detergents used during sacculi isolation (Alvarez et al., 2016; Kuhner et al., 2014). Yet, current protocols are still far from being truly high throughput to be used in large-scale studies.

Here, we have developed a high throughput (HT) method for PG isolation and LC-based analysis appropriate for hundreds to thousands of samples of both Gram-negative and Gram-positive bacteria. This approach has empowered an unprecedented breadth and detail in PG homeostasis genetic determination and PG functional networks. As a proof of concept, we have screened the muropeptide profiles of the non-redundant mutant library of the Gram-negative pathogen *Vibrio cholerae*. *V. cholerae* genome-wide PG profiling revealed a much broader interrelation between PG homeostasis and other cellular processes/pathways than anticipated. Paradoxically, multiple mutants in cell wall genes did not show significant changes in the chemical structure of the PG thus underscoring the existence of functional redundancies and/or conditional determinants. Correlative analyses have allowed to globally visualize the degree of interdependence between PG features as well as potential compensatory mechanisms.

In-depth scrutiny of the *V. cholerae* genome-wide PG profiling uncovered *vc1321.* Despite its annotation as a hypothetical protein and the apparent lack of phenotype, our analysis revealed that VC1321 is a novel high molecular weight penicillin-binding protein (PBP1V, for <u>P</u>enicillin <u>B</u>inding <u>P</u>rotein <u>1</u> of <u>Vibrio</u>) that contributes to *V. cholerae* PG amount and crosslink. Collectively, these results highlight the capability of our innovative HT PG profiling method to perform genome-wide PG analysis to overcome phenotypic and genomic annotation hurdles towards unveiling the full repertoire of cell wall determinants in bacteria.

## RESULTS

### Development of a high throughput method for peptidoglycan isolation

The most commonly used PG isolation protocols were developed in the 60’s (Kolenbrander and Ensign, 1968; Porfirio et al., 2019). These methods capitalize on the separation of SDS-insoluble PG (i.e., murein) from the rest of the cell components and require an individual processing of the samples incompatible with large-scale analyses (**Fig. 1a,b**). First, the bacterial suspension is added dropwise into boiling SDS and stirred for at least one hour to guarantee the complete lysis. Prior to the enzymatic treatment that renders the soluble muropeptides from the macromolecular murein, the detergent is washed away by repeated ultracentrifugation. To overcome these roadblocks, we explored alternative methods compatible with HT sample processing (**Supplementary Fig. 1a**). To this end, we boiled bacterial cells with detergent using different heat transmission systems: water bath, in a heat block (dry) or autoclave. Autoclaving performed the best compared to the other methods, yielding the highest PG amount from the same starting material. Since autoclaving can be easily scaled up to 96-well plates, we selected this approach for the first step.

**Fig. 1.**
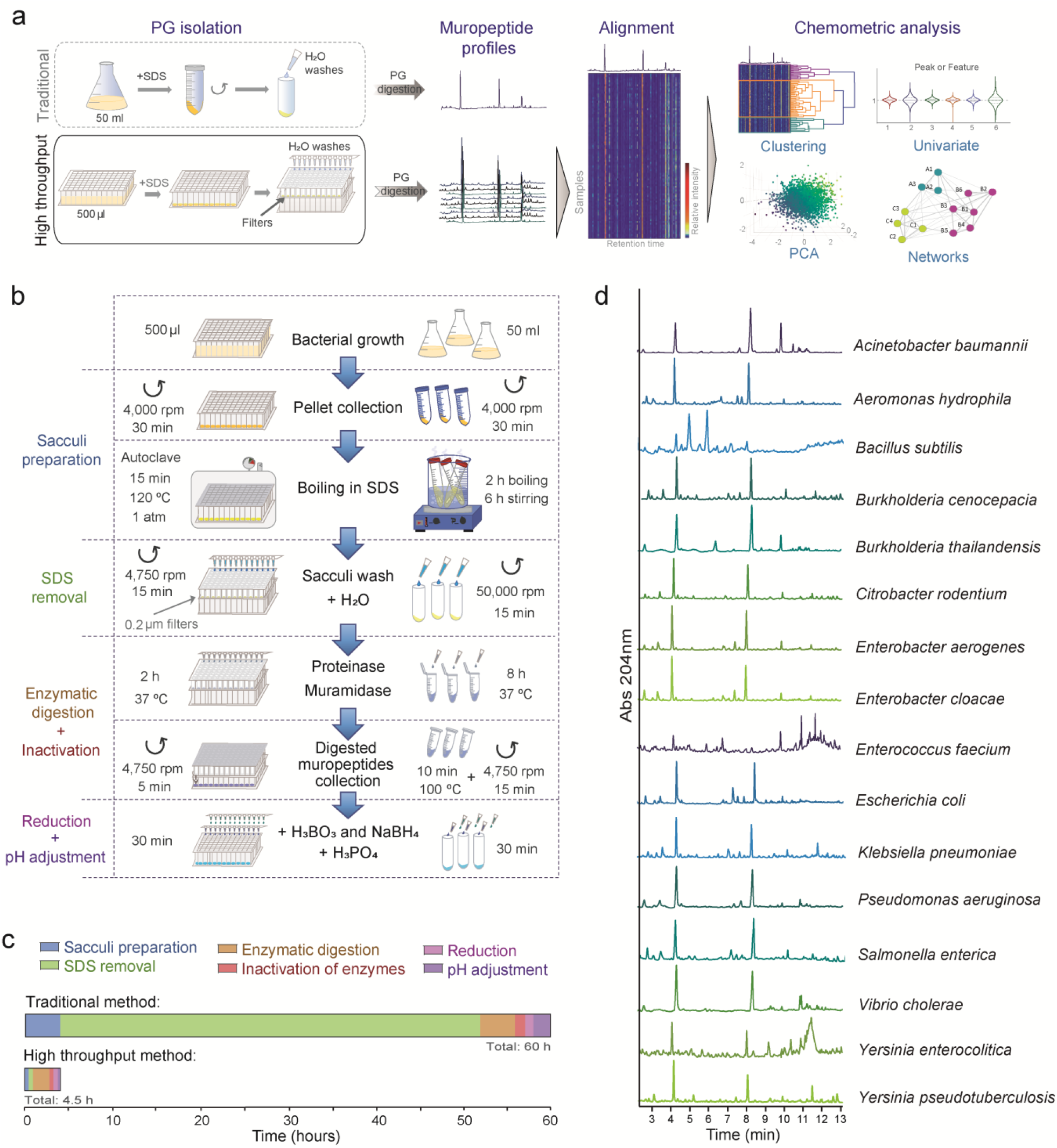
Development of a high throughput method for peptidoglycan sample preparation and analysis. **(a)** Schematic of the comparison between the traditional and the novel high throughput (HT) method for PG isolation and examples of the potential types of analyses that can be carried out with large-scale muropeptide profile datasets. **(b)** Comparative pipeline for the HT and the traditional PG isolation method. **(c)** Comparative timescales for the isolation of PG from 96 samples following the traditional or the HT sample preparation method. **(d)** Representative PG profiles obtained for diverse bacteria using the HT PG isolation method. **See also Supplementary Fig. 1.**

Removing the SDS by series of ultracentrifugation steps is time consuming. Further, since a considerable amount of the sample is lost it requires sufficient cells to produce good quality PG analysis. Traditionally, cultures ranging from 10-100 ml (ca. 10^11^ cells for *E. coli)* are typically used as starting material. This is an obvious bottleneck that limits not just the scale and throughput of the studies, but also the type of samples that can be successfully processed by this protocol, i.e., low-concentrated environmental or *in vivo* samples. As purified sacculus retains the shape of the cell (Formanek and Formanek, 1970) we reasoned that filtration methods used for bacteria could also work for sacculi isolation. Using 0.2 μm pore-size hydrophilic polypropylene (GHP) filters we washed away the SDS from sacculi of different bacteria species (**Supplementary Fig. 1b**) without significant sample loss, thereby allowing the analysis of smaller starting cultures. In fact, SDS-sacculi from only 500 μl of a saturated bacterial culture (~10^9^ total cells) was successfully processed.

Since the sacculi are retained by the filters we reasoned that the subsequent enzymatic digestion steps could be performed in-a-pot. To this end, after protease treatment, we discarded the flow-through and the retained sacculi were subjected to digestion with muramidase to finally elute the soluble muropeptides. Additional modifications of the original protocol were implemented to further cut down the total processing time: i) the final SDS concentration was lowered (from 5 to 1.5% (w/v)) to reduce the number of washes required, ii) the time of the enzymatic reactions was adjusted to ensure their best performance in the shortest time iii) and the reaction buffer was substituted by water since the muramidase performs equally in both solutions (**Supplementary Fig. 1c,d**). Finally, we also adapted for automated liquid handling the sodium borohydride treatment that reduces NAM residues to NA-muraminitol to avoid double peaks due to α- and β-anomeric configurations. All in all, these modifications resulted in the development of an isolation procedure compatible with HT methods using multiwell plates and automated pipetting systems to process simultaneously large number of samples more than ten times faster than the traditional methods (**Fig. 1b,c**). This novel filter-based PG isolation protocol together with the higher speed and sensitivity of UPLC systems (Alvarez et al., 2020; Alvarez et al., 2016) enables the production of large chromatographic datasets for a great range of bacterial species, including both Gram-negative and Gram-positive bacteria (**Fig. 1d**), that can be further analyzed using a variety of bioinformatic tools (Kumar et al., 2017).

### Validation of the high throughput muropeptide profiling

As proof of concept, we used the model organism *V. cholerae.* The muropeptides of the wildtype strain were first identified by UPLC-MS (**Supplementary Fig. 2a,b**). PG analysis of the wildtype strain and over 900 samples grown in duplicates resulted in highly correlated relative abundances for each muropeptide (Pearson correlation score 0.994) (**Fig. 2a**) and demonstrated the robustness and reproducibility of the method. To assess the potential of the HT PG isolation method in discovering novel genetic determinants involved in PG homeostasis, we first analyzed as controls well-characterized *V. cholerae* mutants in cell wall biosynthetic genes: the bifunctional PBP1A (Dorr et al., 2014b), the LD-transpeptidase LdtA (Cava et al., 2011) and the DD-endopeptidase DacA1 (Moll et al., 2015), all of them known to present specific structural changes in their PG. To facilitate comparative analysis between samples, the muropeptide profiles (**Fig. 2b**) were transformed into numeric data. The relative abundance for each muropeptide was calculated by integrating the area under individual peaks divided by the total area of the whole chromatogram. The Log2 fold change (Log2FC) was calculated for every muropeptide using the wildtype values as control (**Supplementary Fig. 2c**). Replicas of each strain properly clustered together and separated from the rest in a principal component analysis (PCA) (**Fig. 2c**). Expected differences between the wildtype PG and that of the three mutant strains were clearly highlighted: the PBP1A^-^ mutant showed a defect in PG amount (**Fig. 2d**); LdtA^-^ a decrease of LD-crosslinked muropeptides, and the DacA1^-^ mutant an increase in pentapeptide-containing muropeptides (**Fig. 2d-f**). Variations in the PG profile of the LdtA- and DacA1 - mutants, correspond to muropeptide changes of ca. 2% of the total area demonstrating that HT PG profiling is reliable to uncover subtle PG variations.

**Fig. 2.**
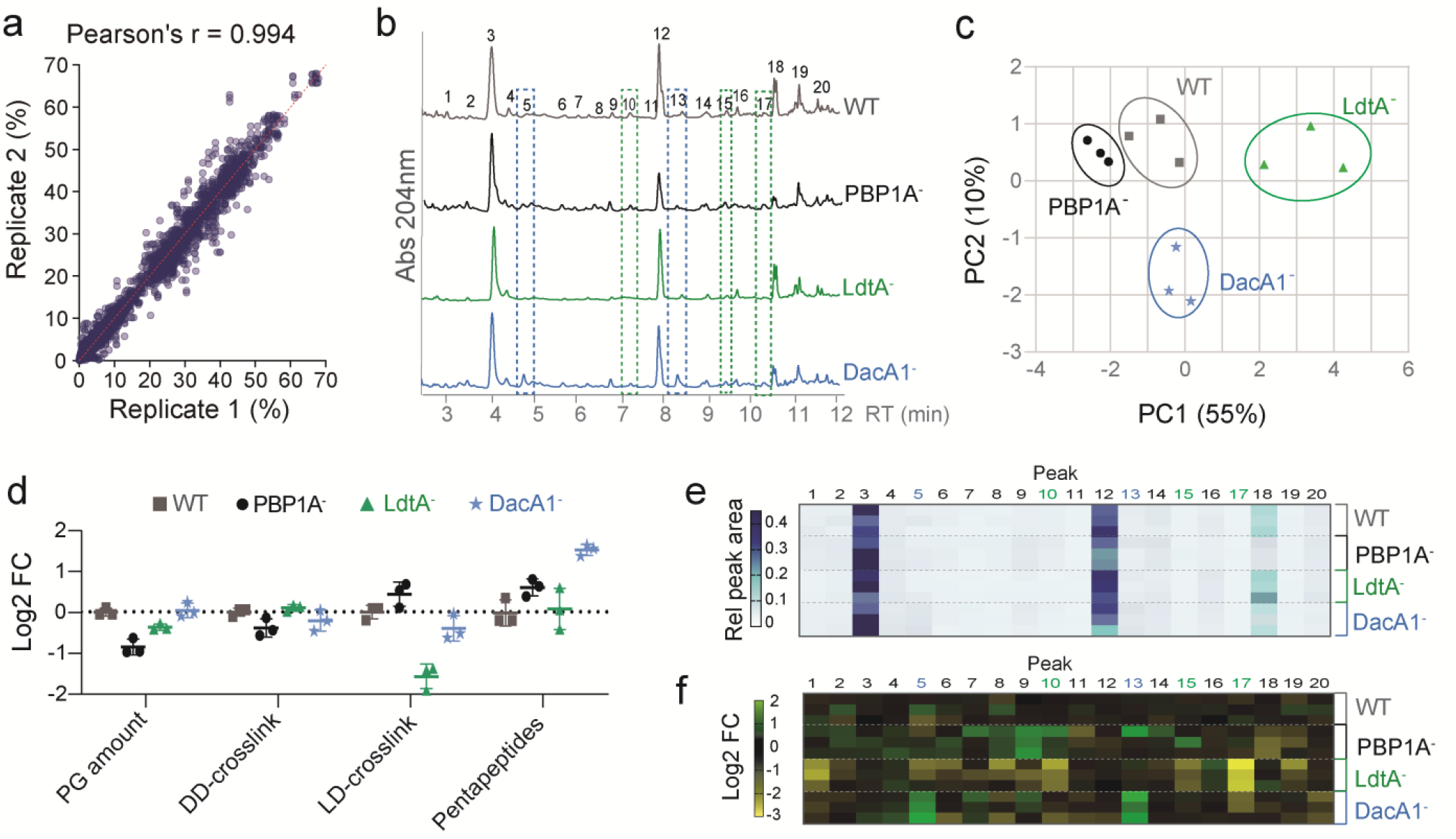
Validation of the high throughput method for peptidoglycan sample preparation. **(a)** Pairwise scatter plot comparing the relative abundance of every muropeptide between replicates of 900 samples. Mean Pearson correlation coefficient is shown. **(b-f)** Validation of the HT method for detection of minor changes in the PG structure by analysis of mutants known to have a phenotype in their muropeptide profile: Pbp1A (PG synthetase), LdtA (LD-transpeptidase) and DacA1 (DD-carboxypeptidase); **(b)** representative chromatogram of each strain; **(c)** PCA using the Log2FC of peak areas of all the muropeptides of each sample; **(d)** analysis of PG features; **(e)** heatmap presenting the relative peak area of each muropeptide numbered in panel **b**; **(f)** heatmap of the Log2FC values (relative to the wildtype) calculated for each peak numbered in **b.** In **e** and **f**, peaks expected to change in LdtA^-^ and DacA1^-^ mutants are highlighted in green and blue respectively. Error bars in **d** correspond to the standard deviation of triplicates. **See also Supplementary Fig. 2.**

### Genome-wide peptidoglycan profiling of *V. cholerae*

To test the potential of the HT PG isolation method in large-scale PG screenings we processed the whole transposon-mutant library of *V. cholerae* (Cameron et al., 2008) (**Fig. 3a**). After discarding inconclusive samples (e.g., mutants that fail to grow), we analysed the PG profile of 3,030 mutants (96% of the library) which altogether generated a comprehensive quantitative dataset for each muropeptide and the main PG-features (**Extended Data 1**).

**Fig. 3.**
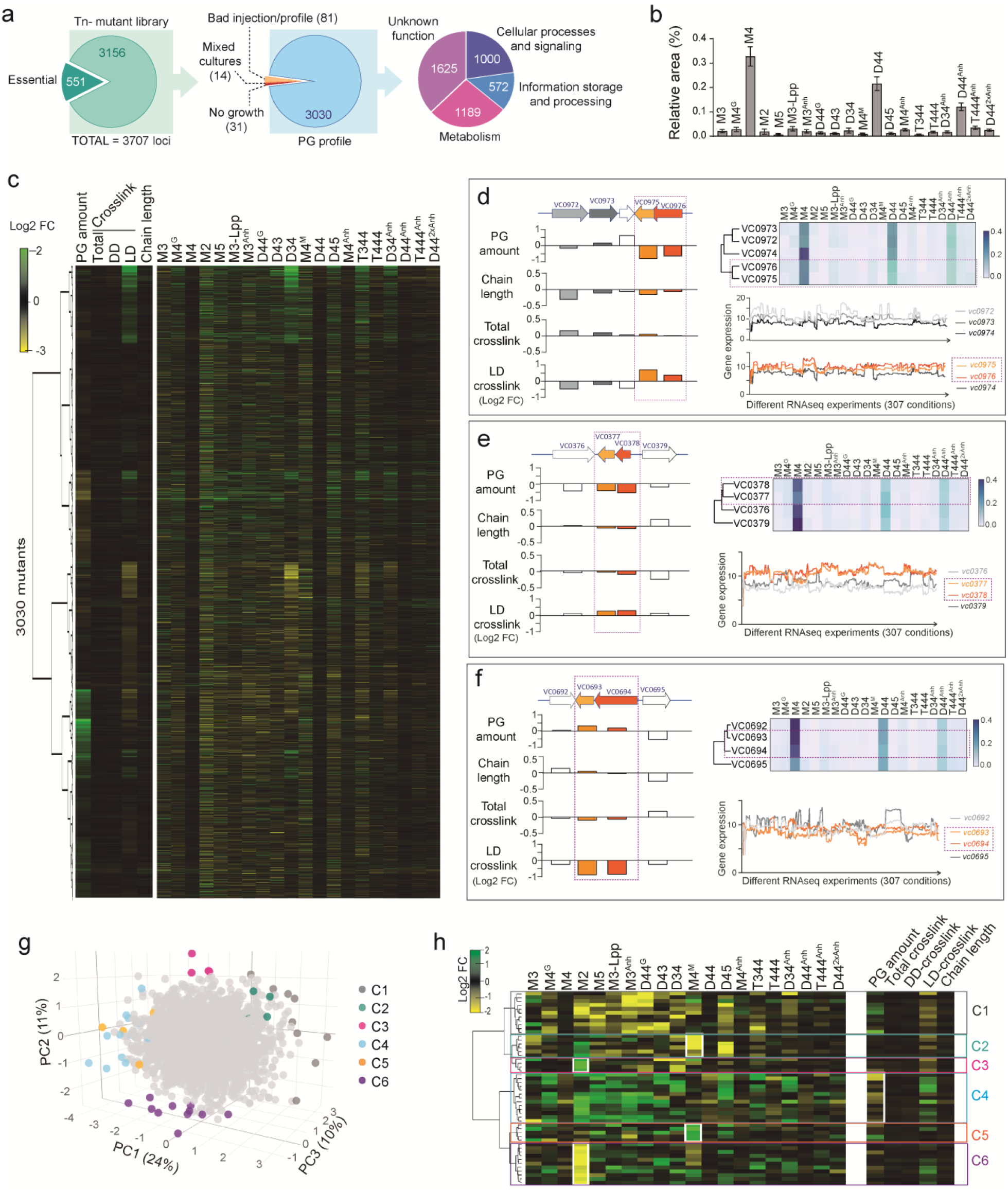
Peptidoglycan profiling of the *V. cholerae* non-redundant mutant library. **(a)** Total number of mutant strains available, processed and analyzed, and their classification into main COG groups. **(b)** Muropeptide abundance and variation (SD) across the whole dataset. **(c)** Hierarchical clustering and heat-map representing the Log2FC calculated for different PG-features and muropeptides. **(d-f)** Analysis of the PG profile of genes belonging to the same transcriptional unit: PG features, heatmap and clustering of the PG profiles of indicated mutants and the expression data of each gene observed in 307 different RNAseq experiments are shown. **(g)** PCA obtained for the mutants of the library based on the Log2FC values observed for the 20 identified muropeptides. Outliers are colour-coded based on their PG-similarities clustering (see **h**). **(h)** Hierarchical clustering of the 50 outliers selected in **g**. **See also Supplementary Fig. 3 and Extended Data 1-3.**

The comparison of the standard deviation (SD) of the relative abundance of each muropeptide in the library dataset with the variation between replicas demonstrated that, except for peaks of very low abundance (i.e., D43 or T344), technical variability is lower than that between biological samples (i.e., mutants), thus supporting the robustness of our method to identify mutants with altered PG (**Supplementary Fig. 3a**). Additionally, we studied the variability of each muropeptide with respect to their relative abundance (**Supplementary Fig. 3b**). Most of the muropeptides adjusted to a linear regression model, except for the M2 and D34 peaks, which show a higher variation degree, indicating that homeostasis of these muropeptides in *V. cholerae* is safeguarded by a high number of genetic determinants.

Relative abundance is not a reliable metric for sample discrimination (**Fig. 3b and Supplementary Fig. 3c-d**). The scree plot and biplot show that the most abundant muropeptides, M4, D44 and D44^Anh^, are the major contributors to the sample variability while fluctuations in less abundant peaks have a more limited impact. On the contrary, Log2FC provided a deeper statistical analysis where the contribution of all peaks had similar weight to define sample’s variability (**Supplementary Fig. 3e**). Hierarchical clustering of the Log2FC values was used to identify sets of genes with similar PG profiles that may belong to the same biological pathway (**Fig. 3c**). We focused on the main PG features: PG amount, type and degree of crosslink, and length of the PG chains. As expected, mutants in genes organized in the same operon (based on multiple RNA-seq datasets) clustered together based on their PG profile (**Fig. 3d-f and Extended Data 2**). As an example, mutation of the loci *vc0975* and *vc0976* results in a significant decrease in the relative amount of PG and an increase in LD-crosslink (**Fig. 3d**). These genes encode two conserved predicted proteases (Teoh et al., 2015) whose function has not been described yet. Similar changes in the PG were observed for the *vc0377* and *vc0378* mutants (**Fig. 3e**), encoding a CheX-like protein proposed to be involved in motility regulation (Altinoglu et al., 2022) and the zinc uptake regulator Zur (Sheng et al., 2015) respectively, which influences PG endopeptidase activity in *V. cholerae* (Murphy et al., 2019). Conversely, mutation of *vc0694* or *vc0693* presented a significant increase of LD-crosslink (**Fig. 3f**). These encode an uncharacterized two component system that has been previously related to intestinal colonization in infant mice (Cheng et al., 2015).

The Log2FC-based PCA for all 20 *V. cholerae* muropeptides revealed outliers representing those mutants with the most defective PGs at a general level (**Fig. 3g**). Even when the PCA is a multiparametric spatial classification of the samples, these outliers grouped in 6 major clusters (**Fig. 3g-h and Extended Data 3**) that shared variability in specific muropeptides (e.g., M2, M4^M^ and D45 muropeptides). The 50 outlier candidates presenting the highest PCA scores included genes of a wide variety of COG (**SI3**) suggesting that multiple cellular pathways preserve bacterial PG homeostasis. As an example, we identified a mutant in *vc0245* (**Fig. 3h**), a gene of unknown function but predicted as an LPS O-antigen polymerase (Manning et al., 1995). This result is in agreement with recent reports pointing the cooperative crosstalk between PG and LPS to maintain envelope integrity (Boll et al., 2016; Goodall et al., 2021; Simpson et al., 2021).

### Genome-wide peptidoglycan profiling reveals correlation between different peptidoglycan features in *V. cholerae*

Multivariate analysis (i.e., PCA) has low resolution to identify genetic determinants of PG homeostasis because while it identifies major general alterations of cell wall’s chemical structure, it fails to discriminate more subtle changes. To solve this bottleneck, we performed univariate data analyses of single PG-features and observed that mutants in known PG-related loci appeared as outliers in the corresponding analyses (**Fig. 4a**). For example, mutation of genes involved in PG synthesis are typically characterized by a defective (sedimentable) PG amount. Accordingly, both the mutant in the PG synthase PBP1A (encoded by *mrcA*), and its cognate regulator CsiV (Dorr et al., 2014a) show up as outliers with low PG amount. Further, the mutant in *vc1718, V. cholerae* homolog of *E. coli*’s envelope biogenesis factor ElyC also displayed low PG amount, in agreement with a previous report (Paradis-Bleau et al., 2014).

**Fig. 4.**
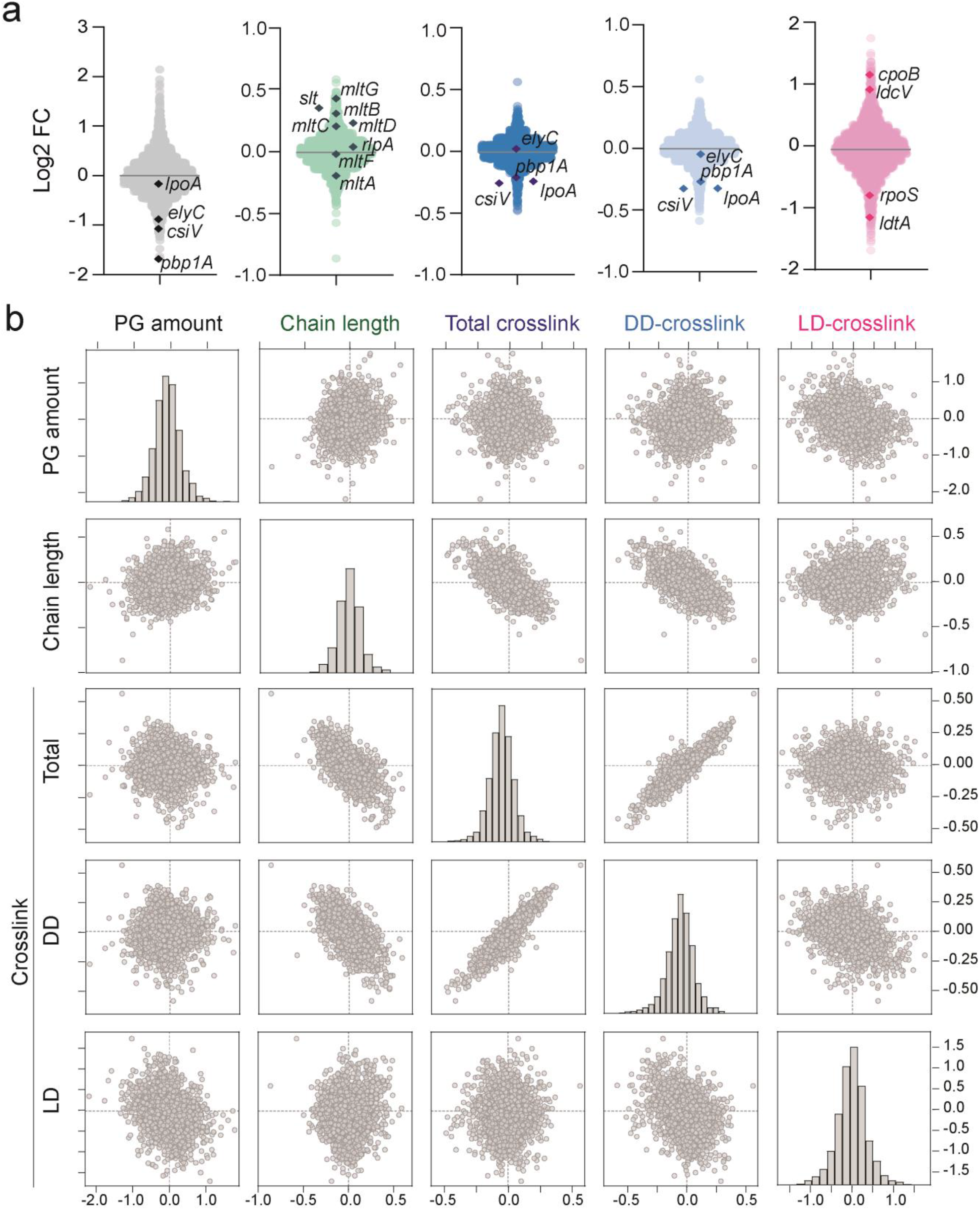
Analysis of the main peptidoglycan features of the *V. cholerae* mutant library. **(a)** Violin plots showing the distribution of samples for the indicated PG-features in the 3030 analyzed mutants. Mutants expected to present a distinctive feature are labelled in each group. **(b)** Correlation scatterplot matrix for the different PG features in the *V. cholerae* mutant library. Histograms showing the distribution of samples for every PG feature are represented in the diagonal.

Additional examples can be found in relation to PG chain length. PG strands terminate in anhydro NAM (Anh) caps generated by the action of lytic transglycosylases (Irazoki et al., 2019). Therefore, quantification of the anhydromuropeptide levels is a proxy to calculate average chain length: high anhydromuropeptide level correlates with shorter PG chains and is indicative of high autolytic activities, while low anhydromuropeptide content is indicative of longer chains typically associated with defects in PG turnover pathways. As expected, most of the mutants in genes encoding lytic transglycosylases presented longer PG chains (**Fig. 4a**). These results are in agreement with recent studies that demonstrated non-redundant roles for some of these lytic transglycosylases in *V. cholerae* (Weaver et al., 2022; Weaver et al., 2019).

The type and degree of the PG crosslinking is another critical aspect of the functionality of the cell wall. *V. cholerae* uses two types of crosslinks: the DD and LD type. DD-crosslinks (AKA 4-3 type) are catalyzed by the DD-transpeptidase activity of certain PBPs such as the bifunctional high molecular weight class A PBPs or the monofunctional class B PBPs. Alternatively, the LD-crosslinks (so called 3-3 type) depend on the activity of LD-transpeptidases (Aliashkevich and Cava, 2021). As for most bacteria, *V. cholerae* primary crosslinking type is DD. This is clearly illustrated in **Fig. 4b** by the direct correlation between DD-crosslink and total crosslink values across mutants. Consistently, mutations in *mrcA* (the bifunctional PBP1A) or its activators *lpoA* or *csiV* resulted in both decreased PG levels and crosslinkage. As mutually inclusive features, PG amount and DD-crosslinking share most of their outliers. However, there are also a number of exceptions (e.g., *elyC* mutant in PG amount but not in crosslinking) thereby suggesting the existence of highly complex functional networks governing PG homeostasis.

In relation with LD-crosslinkage, our data show that the mutant lacking the LD-transpeptidase A (LdtA) and its regulator RpoS are deficient in this crosslinkage type (Cava et al., 2011). Also consistent with recent reports (Hernandez et al., 2020), the mutant in the recycling LD-carboxypeptidase LdcV shows elevated LD-crosslink. Interestingly, a mutant in *vc1834*, which encodes the homolog of *E. coli’s* coordinator of PG synthesis and outer membrane constriction CpoB (Gray et al., 2015), also showed high levels of LD-crosslink. In *E. coli* CpoB is associated to PBP1B, its main PG-synthetase, but in *V. cholerae* PBP1A plays a more prominent role in cell wall synthesis than PBP1B and deletion of PBP1B in *V. cholerae* has not produced any PG phenotype (Dorr et al., 2014b). In *V. cholerae* the two bifunctional PBPs PBP1A and PBP1B have been shown to play distinct roles than in *E. coli*, the PG-phenotype observed here for the CpoB homolog mutant also suggests a different role for this PG-synthesis coordinator in *V. cholerae*, compared to *E. coli.*

Finally, our genome-wide PG analysis also revealed a clear inverse correlation between chain length and total crosslink (**Fig. 4b**). This is consistent with previous observations (Quintela et al., 1995) supporting that a higher number of peptide-crosslinks is required to maintain the mesh integrity of sacculi consisting of shorter glycan chains in average. Collectively these data show that univariate analysis complements the discovery potential of the genome-wide PG profiling.

The normal distribution of the data (**Fig. 4b**) classified as outliers those samples that deviate 1, 2 or 3 SD for at least one muropeptide or PG feature. These data revealed an unexpectedly high genomic contribution to PG homeostasis (**Supplementary Fig. 3g**) that spans beyond traditionally implicated cell wall enzymes and regulators. Interestingly, a Venn diagram representing the outliers that deviate more than 1SD revealed that more than half of the identified mutants (57.4%) displays changes in more than one PG feature. In this line, the aforementioned contribution of DD-crosslink to the total crosslink is supported by the high number of mutants with a phenotype for both features (159), compared to the features alone (27 and 66 respectively). Collectively, genome-wide PG analysis permitted to appreciate a high degree of genomic contribution and cell wall structural interdependence.

### Identification of a new high molecular weight penicillin binding protein in *V. cholerae*

To identify biological processes related with the change of the different PG features, we performed gene-ontology (GO) enrichment analyses using the sets of mutants with significantly altered PG features (i.e., low or high PG amount, anhydro muropeptides, and different kind of crosslink) **(Supplementary Fig. 4 and Extended Data 4)**. We focused on the samples with lower PG amount, since determinants of PG synthesis are a common target for antibiotics. The biological processes “peptidoglycan metabolism” and “peptidoglycan-based cell wall biogenesis” appeared significantly enriched (p-val <0.05) (**Fig. 5a**). In addition to genes encoding known PG-related proteins (e.g., the synthetases PBP1A and PBP1B, or the PG degradative enzymes MltA, MltF and DacA2) (Alvarez et al., 2021), we found *vc1321*, which was annotated as a hypothetical protein in the KEGG database (**Fig. 5b**). We constructed the deletion mutant *Δvc1321* and its complemented strain derivative. Analysis of the PG confirmed that the disruption of this gene not only impairs the PG amount, but also decreases the amount of total crosslink (**Fig. 5c,d**). We confirmed that VC1321 can be specifically labelled by the fluorescent penicillin Bocillin-FL, strongly suggesting that this protein is a new type of class A PBP and so we named it as PBP1V (for Penicillin Binding Protein 1 of Vibrio) (**Fig. 5e**). Sequence similarity and identity was higher between PBP1A and PBP1B than between these PBPs and PBP1V (**Fig. 5f**). *In silico* analysis predicted that PBP1V contains an N-terminal transglycosylase domain (TG) and a putative C-terminal transpeptidase (TP) domain that is split by a region with no homology to previously characterized class A PBPs (**Fig. 5f**). Based on sequence alignments with other high molecular weight PBPs (Sauvage et al., 2008) (**Supplementary Fig. 5**), we selected and mutated three residues: the E172 as the putative TG active site; and S504 and S610 as the potential TP active serines in the SxxK and SxN conserved motifs, respectively (**Fig. 5f**). Mutation of S504 but not of S610 impaired PBP1V binding of Bocillin-FL suggesting that S504 is the TP catalytic serine. Interestingly mutation of E172 also prevented Bocillin-FL binding suggesting a regulatory role of the TG domain in PBP1V transpeptidation activity (**Fig. 5g**).

**Fig. 5.**
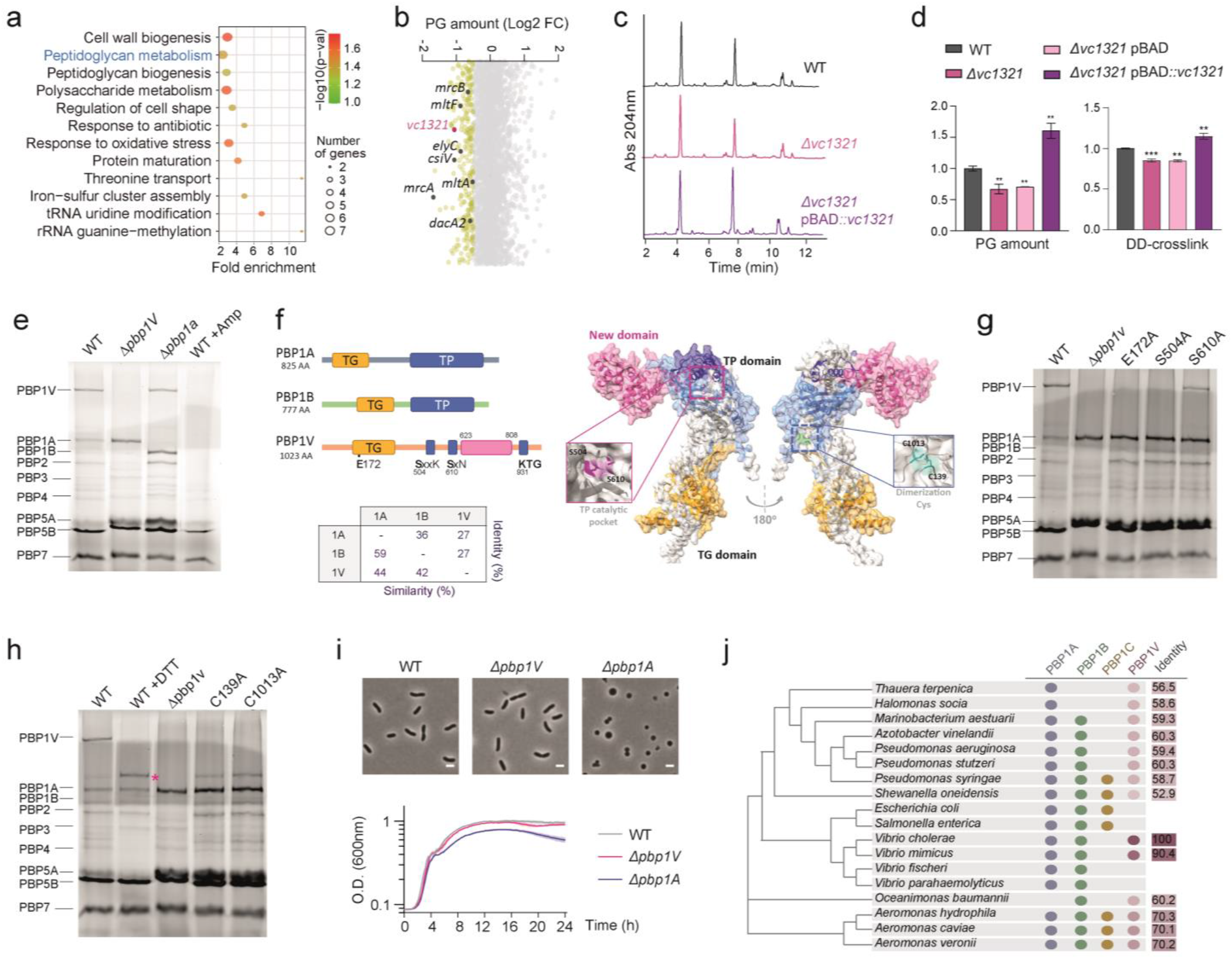
*vc1321* encodes a new high molecular weight PBP in *V. cholerae*. **(a)** Over-represented biological processes associated to those mutants presenting low relative amount of PG (Log2FC <-0.5). The most representative biological processes with a fold enrichment higher than 2 are represented. Dot size indicates the number of loci/category and colour indicates the significance of the enrichment (-Log10(*P*-values)). **(b)** Plot showing the Log2FC of the PG amount for the complete dataset. Genes belonging to the ‘‘peptidoglycan metabolism” category in **a** are labelled. **(c)** Representative PG chromatograms of the wildtype, the *Δvc1321* mutant strain and the *Δvc1321* strain overexpressing *vc1321* from the arabinose-inducible promoter of the pBAD plasmid. **(d)** Quantification of the relative PG amount and DD-crosslink of the wildtype, *Δvc1321* mutant strain, and the *Δvc1321* carrying the pBAD empty plasmid as control or overexpressing *vc1321* from the arabinose-inducible promoter. **(e)** Bocillin gel of indicated strains showing the band corresponding to the product of *vc1321* (named PBP1 of *V. cholerae*, PBP1V) **(f)** Schematic representation of the transglycosylase (TG) and transpeptidase (TP) domains predicted in the sequence of VC1321. **(g-h)** Bocillin gels of *vc1321* point mutants in different putative catalytic residues **(h)** and in candidate cysteines involved in the dimerization of the protein. **(i)** Deletion of *pbp1V* does not result in growth or morphological defect. Scale bar = 2 μm. **(j)** PBP1V homologs across different species. **See also Supplementary Figs. 4-6 and Extended Data 4.**

As PBP1V’s molecular weight was larger than the theoretical value, we hypothesized it could be present in a dimeric form. We noticed that the sequence of PBP1V has two cysteine residues that could be involved in the formation of disulfide bridges between subunits (C139 and C1013). Effectively, *V. cholerae* wildtype membrane fraction treated with the reducing agent DTT resulted in a band shift which coincided with the monomeric size (116 KDa). This result was further confirmed by mutating any of these cysteines (**Fig. 5h**). PG analysis of these mutants demonstrated that the oligomeric state of PVP1V does not affect its role in PG synthesis under the condition tested (**Supplementary Fig. 6**).

Interestingly, unlike the PBP1A mutant, the absence of PBP1V did not impair proper morphology or growth under tested conditions (**Fig. 5i**) reinforcing the idea that genome-wide PG profiling is a powerful tool to identify novel genetic determinants on PG homeostasis, some of which might lack functional annotation or play relevant biological roles in specific conditions. Finally, while PBP1V is only conserved in *V. cholerae* and its close relative *V. mimicus* within the genus of *Vibrio*, orthologs of this PBP are also encoded by other species frequently associated with marine lifestyles (**Fig. 5j**).

## DISCUSSION

We have developed a filtration-based method for PG isolation adapted to 96-well plates and automated pipetting systems that enables HT PG biochemical analyses. We have proved that the new protocol is applicable to different bacterial species and provides PG profiles comparable to those obtained using the traditional procedure. The use of filters drastically reduces the initial number of bacteria required for PG isolation by ca. 2 orders of magnitude compared to the minimum amount required by protocols using ultracentrifugation. Therefore, in addition to enabling larger-scale analyses, this method opens up the possibility to characterize the PG chemical structure of low-PG challenging samples such as intracellular non-proliferating bacteria. Moreover, as filtration prevents PG loss, the limiting factor with this method is no longer the sample size (amount of bacteria) but rather the detection limit of the analytical system. In this regard, the accuracy and sensitivity of current MS has been improving during the last decades and can facilitate the muropeptide characterization of challenging low-abundant PG samples.

Developing a HT PG isolation method has been instrumental to analyse the cell wall of the non-redundant mutant library of *V. cholerae*, i.e., the first genome-wide PG profiling of a bacterium. Analysis of the data confirmed the role of known PG-related genes and established new links between previously unrelated genes with cell wall homeostasis, including genes encoding for proteins with previously described or unknown functions. The unprecedented width of the omic-scale PG profiling has made possible to witness a remarkably high genomic contribution to PG homeostasis. For instance, almost 98% of the mutants of the *V. cholerae* transposon library present a statistically significant alteration of their PG profile (deviate at least 1SD from the average), and more than the 50% of the mutants presented a variation higher than 2SD of the average of the whole library for at least one muropeptide or a PG-feature) that included loci from all COG categories. These results are in agreement with studies that linked the cell wall with seemingly unrelated pathways/processes (e.g.,(Bancroft et al., 2020; Campbell et al., 2021; Masser et al., 2021)). Conversely, certain mutants in cell wall-annotated genes did not show an altered PG profile suggesting the existence of functional redundancies or condition-specific activities. Future research should uncover the functional relationships within these complex genetic networks and what regulatory rewiring occurs in response to environmental changes.

Additionally, we observed that variations in PG features and muropeptides are in general mutually inclusive: i.e., a significant proportion of the mutants present changes in more than one PG feature. This result is consistent with the idea of PG structure being safeguarded by compensatory mechanisms that preserve the cell wall integrity. This balance can be explained following a “substrate-product” logic as it is for the inverse relationship between monomers and dimers or between pentapeptides and tetrapeptides, but often seems to be the result of more complex regulatory associations (e.g., crosslink vs chain length) implicating multiple variables.

A main strength of this global study is its potential to identify novel cell wall genetic determinants even when lacking a growth phenotype under the conditions studied. As an example, we discovered PBP1V, a third high molecular weight PBP in *V. cholerae* which differs from the previously characterized PBP1A and PBP1B of this bacterium and is also not homolog to PBP1C proteins found in other bacteria. Deletion of *pbp1V* did not produce any growth defect in our analyses, but it decreased the total amount and the degree of crosslink of the PG. Although only the TG domain was predicted for this protein, sequence alignment with other high molecular weight PBPs suggested a potential TP domain consisting of the canonical active site sequences separated by a 186 amino acids domain that was not previously described in any bi-functional PBPs so far studied. Mutation of the S504 residue of PBP1V that forms part of a conserved S*xxK TP motif, prevented binding of PBP1V to Bocillin FL, strongly suggesting that this is the catalytic serine of the TP domain. Surprisingly, mutation of E172, which is not found into a conserved EDxxFxxHxG sequence typical of TG domains, also affected binding of Bocillin FL, suggesting that the TG activity might also influence the TP activity of PBP1V’s transpeptidation, similar as previously reported (Zijderveld et al., 1991). PBP1V was found to form dimers by intermolecular disulfide bonds between cysteines C139 and C1013. Dimerization of high molecular weight PBPs by disulfide bonds has been previously reported as a mechanism of redox control of their activity (Bukowska-Faniband and Hederstedt, 2017). However, the monomeric PBP1V retains its TP activity suggesting that dimerization might rather affect other aspects such as localization or protein interactions.

The HT method presented here for PG isolation and genome-wide PG profiling represents a promising tool for discovering new genetic determinants of PG biosynthesis and remodeling (i.e., PBP1V) and their genetic networks, the functional relations between the PG features and the structural constraints that define a “healthy” PG sacculus. By profiling mutant libraries of diverse bacterial species (including some of the most important bacterial pathogens), the publicly available experimental platforms and databases generated will have an even wider impact due to their utility to uncover general and species-specific mechanisms underpinning PG homeostasis. Additionally, by using diverse growth conditions we can learn how bacteria adapt their cell wall to relevant environments (e.g. the host) and which constraints cannot be surpassed in the PG structure to preserve viability and shape. Interlinking this PG-biochemical information with phenotypic analyses of the same mutant libraries (Nichols et al., 2011) will permit to create highly informative PG profile-phenotype-gene clusters to better understand the biological consequences of PG synthesis and remodeling. Finally, since the cell wall is one of the major “Achilles heels” of bacteria, this systematic survey of new players in the synthesis and regulation of the PG structure will be particularly relevant on antibiotic resistant pathogenic bacteria and infection-mimicking conditions as it has a great potential to promote the discovery of new pathways that can be targeted in antimicrobial therapies.

## Supporting information

Extended Data 2 - SRA Accessions

Extended data 3 - PCA Outliers

Extended Data 4 - Enrichments

Extended Data 5 - Bacterial strains

Extended Data 6 - Primers

## ACKNOWLEDGEMENTS

Research in the Cava lab is supported by The Swedish Research Council (VR), The Knut and Alice Wallenberg Foundation (KAW), The Laboratory of Molecular Infection Medicine Sweden (MIMS) and The Kempe Foundation. We thank John J. Mekalanos for providing the *V. cholerae* transposon mutant library and the Cava Lab and Miguel A. de Pedro for their helpful discussions.

## AUTHOR CONTRIBUTION

SBH, LAM and FC designed the experiments. SBH, LAM and BRR performed research. SBH, LAM, BRR, BS and AS analysed data. SBH, LAM and FC wrote the paper with input from the rest of the authors.

## SUPPLEMENTARY FIGURES TITLES AND LEGENDS

**Supplementary Fig. 1. High throughput method optimization, Related to Fig. 1.**

**(a)** Comparison of the efficiency of PG isolation methods. Samples corresponding to different cell numbers were collected, supplemented with SDS 1.5% (w/v) final concentration, and solubilized by boiling, autoclaving or using a heat-block during different times. Samples were processed and analyzed by UPLC. Yield is quantified as maximum intensity of absorbance at 204 nm for the M4 muropeptide, in arbitrary units (AU). **(b)** Phase contrast microscopy of different bacteria and their purified sacculi processed using the traditional or the HT purification method. Scale bar = 2 μm. **(c)** Relative muramidase activity at different time points. **(d)** Efficiency of the muramidase digestion in buffer or water on sacculi from two different concentrated starting cultures at two time points. Yield is quantified as maximum intensity of absorbance at 204 nm for the M4 muropeptide, in arbitrary units (AU).

**Supplementary Fig. 2. Muropeptide profile of *V. cholerae*’s peptidoglycan obtained using the high throughput sample preparation method and analysis of data, Related to Figs. 2-3.**

**(a)** Representative chromatogram obtained for *V. cholerae* wildtype strain using the HT sample preparation method. Peaks of interest are listed. **(b)** Table of identified muropeptides in **a**. Identity was confirmed by MS/MS analysis, theoretical and observed neutral mass in Da are indicated. GlcNAc: N-acetyl glucosamine; MurNAc: N-acetyl muramic acid; (1-6anhydro) MurNAc: 1-6 anhydro N-acetyl muramic acid, terminal muropeptide; L-Ala: L-alanine; D-Glu: D-glutamic acid; DAP: *meso*-diaminopimelic acid; D-Ala: D-alanine; D-Met: D-methionine; Gly: glycine; Lpp: Braun’s lipoprotein. **(c)** Pipeline for the data transformation and analysis: i) abundance was calculated by integration of the area of the peaks, ii) relative area of each muropeptide was calculated by dividing the peak area by the total area of the chromatogram, iii) finally, fold change (FC) relative to the control sample was calculated and Log2FC was used for representation in heat maps.

**Supplementary Fig. 3. Analysis of the peptidoglycan profiling of the *V. cholerae* non-redundant mutant library, Related to Fig. 3.**

**(a**) Biological versus technical variability: comparison of the standard deviation of the relative abundance of each muropeptide across all samples in the *V. cholerae* mutant library versus the average standard deviation across replicas of 300 samples. **(b)** Correlation between each muropeptide relative abundance and its standard deviation across the library dataset.

**(c)** Scree plot of the main components used for the PCA using relative abundance of muropeptides and Log2FC calculated values. **(d)** Biplot showing the contribution of the different variables (muropeptides) to the data variability in the dataset of relative amounts**. (e)** Biplot showing the contribution of the different variables (muropeptides) to the data variability in the Log2FC dataset. **(f)** Violin plots showing the distribution of the samples for each muropeptide. Log2FC data was used. Median (continued lines) and quartiles (dashed lines) are represented. **(g)** Table representing the number and percentage of mutants with a significant phenotype for at least one muropeptide or a PG feature. A significant phenotype is considered when the sample is further than 1, 2 or 3 standard deviations (SD) from the average. **(h)** Venn diagram showing the number of outliers with a phenotype (using 1SD as threshold) in the main PG features.

**Supplementary Fig. 4. Enrichment analysis of mutants presenting differential PG features, Related to Fig. 5.**

Over-represented biological processes of mutant candidates presenting low (left, red dots) and high (right, blue dots) relative amounts of: **(a)** chain length (Log2FC <-0.2 and >0.18); **(b)** total crosslink (Log2FC <-0.15 and >0.15); **(c)** DD-crosslink (Log2FC <-0.2 and >0.2); and **(d)** LD-crosslink (Log2FC <-0.7 and >0.7). The most representative biological processes with a fold enrichment higher than 2 are represented. The dot size in the balloon plots indicates the number of loci included in each category and the colour the significance of the enrichment (-Log10(p-values)).

**Supplementary Fig. 5. Multiple sequence alignment of PBP1V with other high molecular weight penicillin-binding proteins, Related to Fig. 5.**

Sequence alignment of *V. cholerae* PBP1V (Vc-PBP1V, Uniprot: Q9KSD7), *V. cholerae* PBP1A (Vc-PBP1A, Uniprot: Q9KNU5), *V. cholerae* PBP1B (Vc-PBP1B, Uniprot: Q9KUC0), *E. coli* PBP1A (Ec-PBP1A, Uniprot: P02918), *E. coli* PBP1B (Ec-PBP1B, Uniprot: P02919) and *E. coli* PBP1C (Ec-PBP1C, Uniprot: A0A093EN65), performed with T-COFFEE Expresso. Background residue colour indicates degree of conservation. TG domain is highlighted in yellow, TP domain is highlighted in light blue, PBP1C-binding domain is highlighted in green. PBPs conserved motifs are indicated in the red boxes. Vc-PBP1V new domain in red letters.

**Supplementary Fig. 6. Peptidoglycan analysis of PBP1V cysteine point mutants**, **Related to Fig. 5.**

Quantification of the relative PG amount and DD-crosslink of the wildtype, *Δpbp1V* mutant strain, and indicated cysteine mutants. (ns: not significant).

## EXTENDED DATA FILES AND LEGENDS

**Extended Data 1.** PG dataset from the *V. cholerae* Tn-library, **Related to Fig. 3.**

**Extended Data 2.** SRA numbers used for the study of gene expression analysis, **Related to Fig. 3 d-f.**

**Extended Data 3.** Data 50 selected PCA outliers, **Related to Fig. 3 g-h.**

**Extended Data 4.** GO-enrichments, **Related to Fig. 5.**

**Extended Data 5.** Bacterial strains list, **Related to Methods.**

**Extended Data 6.** List of primers, **Related to Methods.**

## METHODS

### Lead Contact and Materials Availability

- All the data obtained from the genome-wide HTPGP of *V. cholerae* non-redundant mutant collection is available in the Extended Data files.
- Further information and requests for resources and data should be directed and will be fulfilled by the Lead Contact, Felipe Cava.

### Bacterial strains, media, and antibiotics

All the bacterial strains used in this study are listed in **Extended Data 5**. Lysogeny broth (LB) containing 10 or 5 mg/ml of NaCl (for *V. cholerae* or the other bacterial species respectively) was used as the standard growth medium. Agar 1.5% (w/v) was used in solid plates. Unless otherwise specified, antibiotics were used at the following concentrations: streptomycin 200 μg/ml, kanamycin 50 μg/ml.

Plasmids were constructed by standard DNA cloning techniques. *V. cholerae* mutant strains were constructed using the primers listed in **Extended Data 6,** and standard allele exchange techniques with derivatives of the suicide plasmid pCVD442 as described previously (Donnenberg and Kaper, 1991). Plasmid used for complementation are derivatives of pBAD18 (Guzman et al., 1995), where expression is controlled by the P_BAD_ promoter. The fidelity of the DNA regions generated by PCR amplification was confirmed by DNA sequencing.

### Growth conditions

Bacterial species were grown in LB at 30 or 37 °C for sacculi isolation. The culture volume used for high throughput sacculi preparation ranged between 0.5 and 2 ml and was adjusted to obtain the best PG-profile (highest absorbance, UPLC signal) without saturating the filters during the purification. For the analysis of the *V. cholerae* mutant library, cells from glycerol stocks were first inoculated in 96-well plates containing 200 μl of LB medium, grown overnight at 37 °C and further diluted 1000-fold into 500 μl of fresh medium in 96-well deep-well plates, where they were finally grown at 37 °C with shaking overnight. For induction of the P_BAD_ promoter, 0.2% (w/v) of arabinose (final concentration) was added to bacterial cultures.

### High Throughput (HT) sacculi preparation and PG isolation and digestion

For optimization of the method, different sacculi isolation protocols were tested, shown in **Supplementary Fig. 1**. Bacterial cultures grown overnight in 96-well deepwell plates were pelleted in the same culture plates at 5,250 *x g* for 30 min. The supernatant was discarded, and the pellets were resuspended with 50 μl of fresh medium. After addition of 25 μl of 5% SDS, samples were tightly sealed using silicone mats and tape, and autoclaved (15 min at 120 °C 1atm). Samples were then transferred to 96-well-filter plates (AcroPrep Advance 96-well 0.2 μm GHP membrane filter plates, 2 ml volume, PALL). SDS was removed by washing twice with 1 ml Milli-Q water (5,250 *x g* for 15 min). SDS-free sacculi samples were treated with 200 μl proteinase K (40 μg/ml in 100 mM TrisHCl pH 8.0, 30 min at 37°C) for removal of the Braun’s lipoprotein. After an additional 1 ml wash with water, samples were digested with 50 μl muramidase (100 μg/ml in water, overnight at 37°C). Enzymatic reactions were performed directly on the filter, sealing the plates with aluminium foil and using a humidity chamber to reduce evaporation.

After muramidase digestion, soluble muropeptides were eluted into a new 96-well deep-well plate by centrifuging the filter plate at 5,250 *x g* for 5 min. Finally, eluted muropeptides were reduced using 10 μl borate buffer (0.5 M pH 9.0) and 25 μl NaBH_4_ (10 mg/ml) for 30 min at room temperature. The pH of the samples was adjusted to 3.5 using 8 μl orthophosphoric acid 25% (v/v) and the samples were stored frozen at −20 °C until analysis by UPLC.

### Data acquisition by liquid chromatography

PG samples were analyzed by liquid chromatography, as previously described (Alvarez et al., 2020). Briefly, muropeptides were separated using an UPLC system (Waters) equipped with a trapping cartridge precolumn (SecurityGuard ULTRA Cartridge UHPLC C18 2.1 mm, Phenomenex) and an analytical column (Waters ACQUITY UPLC BEH C18, 130Å, 1.7 μm, 2.1 mm×150 mm), using as solvents 0.1% formic acid in Milli-Q water (buffer A) and 0.1% formic acid (v/v) in acetonitrile (buffer B) in a 14 min run. Muropeptides were detected by measuring the absorbance at 204 nm. Identity of the peaks was first assigned by LC coupled to a QTOF-MS instrument and then by comparison of their retention time. The QTOF–MS instrument was operated in positive ionization mode. Detection of muropeptides was performed by MS^E^ to allow for the acquisition of precursor and product ion data simultaneously, using the following parameters: capillary voltage at 3.0 kV, source temperature to 120 °C, desolvation temperature to 350 °C, sample cone voltage to 40 V, cone gas flow 100 l/h, desolvation gas flow 500 l/h and collision energy (CE): low CE: 6 eV and high CE ramp: 15-40 eV. Mass spectra were acquired at a speed of 0.25 s/scan. The scan was in a range of 100-2000 *m/z.* Data acquisition and processing was performed using UNIFI software package (Waters Corp.).

For the study of the PG chemical variability between samples, collected UPLC chromatographic data were analyzed using the PG-metrics pipeline (Kumar et al., 2017). In brief, raw data were pre-processed by trimming out irrelevant segments and by baseline correction subtracting a synthetically created baseline. Trimmed and baseline corrected datasets were then aligned by segments using the COW algorithm by selecting appropriate reference samples and combinations of the segment length and slack parameters. The peak area of aligned chromatograms was calculated using a MATLAB routine based on the trapezoidal integration approach wherein the boundaries of the integration were manually selected.

The relative area of each muropeptide was calculated dividing its peak area by the total area of the chromatogram (**Supplementary Fig. 2b**). The different PG features were calculated as described previously (Alvarez et al., 2020; Vigouroux et al., 2020). When relative peak areas of each mutant were used, the most abundant muropeptides presented the greatest impact on the results of PCA analyses (**Supplementary Fig. 3a-c**), thus the log2 fold change (Log2FC) was calculated for more comprehensive analysis of the data. Log2FCs were calculated by dividing each value (peak area or PG-feature) by the one of the control sample when available or by the mean value of all samples from the same sample set (i.e., 96-well plates) (**Supplementary Fig. 2c**).

### Data analysis and representation

Prism 8.0 (GraphPad Software) was used to plot and analyze numerical data. Statistical tests in Prism calculated significance of measurements as reported in the corresponding figure legends.

Statistical analyses (correlation, standard deviation, PCA, biplots, scree plots) were carried out using R version 4.1.2.

Venn diagrams were created selecting as outliers those mutants with a PG phenotype (more than 1, 2 or 3 standard deviations) in at least one muropeptide or PG feature.

Heatmaps and hierarchical clustering analyses (agglomeration method: ward; distance metric: euclidean) were performed using the NG-CHM Heat Map Builder, version 2.20.2 (Ryan et al., 2019).

### Gene expression analysis

In order to calculate gene expression estimates, 307 RNA-Sequencing data annotated to *V. cholerae* and performed on an Illumina sequencing machine were downloaded from the NCBI’s sequence read archive (SRA) (Leinonen et al., 2011) (See **Extended Data 2**). We then used salmon/v1.3.0 (Patro et al., 2017) with parameters --libType A --seqBias and --posBias to quantify expression against a decoy-aware index generated from the GCF_000829215.1_ASM82921v1 reference (Okada et al., 2014) as described in Srivastava et al. (Srivastava et al., 2020). Post quantification, we filtered experiments where less than 50% of reads could be matched to a reference gene and imported the estimated counts into R/v4.0 using the tximport package (Soneson et al., 2015). After importing, we used the varianceStabilizingTransformation procedure implemented in the DESeq2 (Love et al., 2014) R package to generate a blind, homoscedastic, and pseudo-log2 count set.

### Functional enrichment analysis

The Protein ANalysis THrough Evolutionary Relationships (PANTHER) Classification System and the PANTHER Statistical Overrepresentation Test (released 20200407) were used to categorize the selected datasets by Gene Ontology (GO) and to determine the overrepresented GO terms (Mi et al., 2019a; Mi et al., 2019b). For this, selected datasets were searched using the “GO biological process complete” annotation classification available in PANTHER to find functional classes statistically overrepresented when compared to the reference list containing the 3030 proteins corresponding to the Tn-mutants analysed by HT-PG profiling. GO terms were considered significantly overrepresented only when the P-value (Fisher’s exact test) was lower than 0.05. The resulting list for each dataset was summarized (similar terms including the same list of genes were removed to avoid redundancy) and visually represented as described by Bonnot *et al.* (Bonnot et al., 2019).

### Microscopy imaging

Bacteria were immobilized on LB pads containing 1% agarose (w/v). Phase contrast microscopy was performed using a Zeiss Axio Imager .Z2 microscope (Zeiss, Germany) equipped with a Plan-Apochromat 63X phase contrast objective lens and an ORCA-Flash 4.0 LT digital CMOS camera (Hamamatsu Photonics, Japan), using the Zeiss Zen Blue software. Image analysis and processing were performed using ImageJ software (Rasband, W.S., ImageJ, U. S. National Institutes of Health, Bethesda, Maryland, USA, https://imagej.nih.gov/ij/, 1997-2018).

### Membrane preparation and detection of penicillin-binding proteins

For the preparation of membranes of *V. cholerae*, 100 ml cell cultures were grown over night and harvested by centrifugation at 15,000 *x g* for 15 min at 4 °C (Beckman JLA-16.250 rotor, Beckman Avanti J-25, Beckman Coulter). Pellets were washed with 20 mM potassium phosphate (pH 7.5) and 140 mM sodium chloride buffer. Following another centrifugation for 30 min at 360 *x g* at 4 °C, cells were disrupted twice using a French press (Pressure Cell Homogenizer FPG12800, Stansted Fluid Power Ltd). The resulting cell lysates were subjected to centrifugation at 360 *x g* for 10 min at 4 °C. The supernatant fractions were collected and centrifuged at 75,600 *x g* for 90 min at 4 °C (Beckman JA25.50 rotor, Beckman Avanati J-25, Beckman Coulter). The resulting pellets were washed again and resuspended in 500 μl potassium phosphate buffer. The concentration of the membrane preparations was measured using Bradford reagent (Bio-Rad) with bovine serum albumin as standard (Sigma).

For the detection of the PBPs, membranes were incubated for 30 min at 37 °C with a fluorescent labelling agent, Bocillin FL Penicillin (Invitrogen, Thermo Fisher Scientific) at a final concentration of 0.05 mM. Finally, the samples were denatured with 4x Laemmli sample buffer (Bio-Rad) at 95 °C for 5 min, followed by centrifugation to remove membrane debris. 20 μl of the reaction mixture were subjected to SDS-PAGE analysis on an 8% polyacrylamide gel. Penicillin-binding proteins were visualised with a laser scanner, Typhoon FLA 9500 (GE Healthcare Life Sciences) at 473 nm with a 530DF20 emission filter.

### Multiple sequence alignments

Sequence from *V. cholerae* PBP1V (Vc-PBP1V, Uniprot: Q9KSD7), *V. cholerae* PBP1A (Vc-PBP1A, Uniprot: Q9KNU5), *V. cholerae* PBP1B (Vc-PBP1B, Uniprot: Q9KUC0), *E. coli* PBP1A (Ec-PBP1A, Uniprot: P02918), *E. coli* PBP1B (Ec-PBP1B, Uniprot: P02919) and *E. coli* PBP1C (Ec-PBP1C, Uniprot: A0A093EN65) were aligned using T-COFFEE Expresso Version 11.00 (Notredame et al., 2000) and visualized with JalView (Waterhouse et al., 2009).

We run BLAST to study the homology between *V. cholerae* HMW-PBPs (Altschul et al., 1997).

### Protein structure prediction

PBP1V protein 3D structural model was built with Alphafold, using both monomer and multimer modes (Evans et al., 2022; Jumper et al., 2021). Only the best model among the five best given by default was examined in detail and represented in the figures. Molecular graphics and analyses were performed with ChimeraX (Goddard et al., 2018; Pettersen et al., 2021).

