## Supplementary Figures for "Genome-wide peptidoglycan profiling of *Vibrio cholerae*"

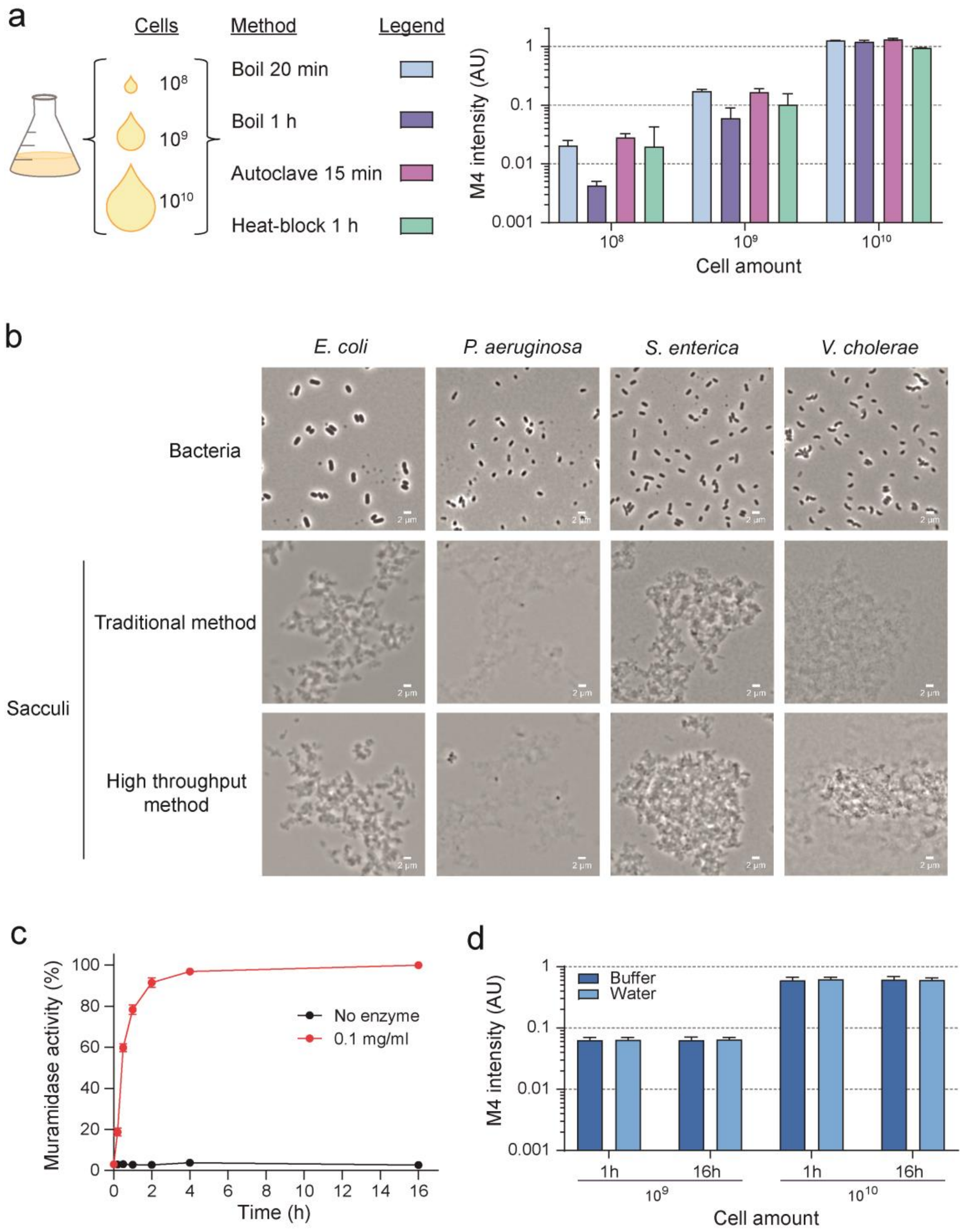


**Supplementary Fig. 1. High throughput method optimization, Related to Fig. 1.**
**(a)** Comparison of the efficiency of PG isolation methods. Samples corresponding to different cell numbers were collected, supplemented with SDS 1.5% (v/v) final concentration, and solubilized by boiling, autoclaving or using a heat-block during different times. Samples were processed and analyzed by UPLC. Yield is quantified as maximum intensity of absorbance at 204 nm for the M4 muropeptide, in arbitrary units (AU). **(b)** Phase contrast microscopy of different bacteria and their purified sacculi processed using the traditional or the HT purification method. Scale bar = 2 µm. **(c)** Relative muramidase activity at different time points. **(d)** Efficiency of the muramidase digestion in buffer or water on sacculi from two different concentrated starting cultures at two time points. Yield is quantified as maximum intensity of absorbance at 204 nm for the M4 muropeptide, in arbitrary units (AU).

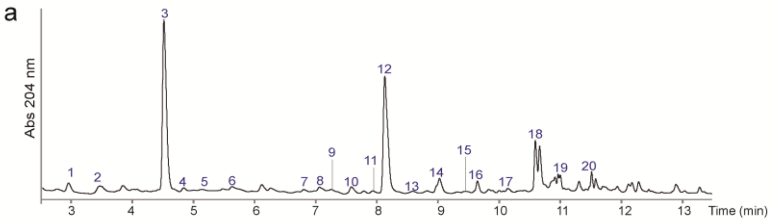


b

| Peak | Name | Structure | Neutral mass (Da) Theoretical | Neutral mass (Da) Observed | Difference |
|---|---|---|---|---|---|
| 1 | M3 | GlcNAc-MurNAc-L-Ala-D-Glu-DAP | 870.3706 | 870.3728 | -0.0022 |
| 2 | M4$^{G}$ | GlcNAc-MurNAc-L-Ala-D-Glu-DAP-Gly | 927.3921 | 927.3908 | 0.0013 |
| 3 | M4 | GlcNAc-MurNAc-L-Ala-D-Glu-DAP-D-Ala | 941.4077 | 941.4080 | -0.0003 |
| 4 | M2 | GlcNAc-MurNAc-L-Ala-D-Glu | 698.2858 | 698.2846 | 0.0012 |
| 5 | M5 | GlcNAc-MurNAc-L-Ala-D-Glu-DAP-D-Ala-D-Ala | 1012.4448 | 1012.4524 | -0.0076 |
| 6 | M3-Lpp | GlcNAc-MurNAc-L-Ala-D-Glu-DAP-Braun's lipoprotein (TK residues) | 1099.5132 | 1099.5144 | -0.0012 |
| 7 | M3$^{Anh}$ | GlcNAc-(1-6anhydro)MurNAc-L-Ala-D-Glu-DAP | 850.3444 | 850.3437 | 0.0007 |
| 8 | D44$^{G}$ | M4-M4$^{G}$ (crosslinked at D-Ala-DAP) | 1850.7892 | 1850.7793 | 0.0099 |
| 9 | D43 | M4-M3 (crosslinked at D-Ala-DAP) | 1793.7677 | 1793.7611 | 0.0066 |
| 10 | D34 | M3-M4 (crosslinked at DAP-DAP) | 1793.7677 | 1793.7593 | 0.0084 |
| 11 | M4$^{M}$ | GlcNAc-MurNAc-L-Ala-D-Glu-DAP-D-Met | 1001.4111 | 1001.4109 | 0.0002 |
| 12 | D44 | M4-M4 (crosslinked at D-Ala-DAP) | 1864.8048 | 1864.8014 | 0.0034 |
| 13 | D45 | M4-M5 (crosslinked at D-Ala-DAP) | 1935.8420 | 1935.8563 | -0.0143 |
| 14 | M4$^{Anh}$ | GlcNAc-(1-6anhydro)MurNAc-L-Ala-D-Glu-DAP-D-Ala | 921.3815 | 921.3804 | 0.0011 |
| 15 | T344 | M3-M4-M4 (crosslinked at DAP-DAP and D-Ala-DAP) | 2717.1649 | 2717.1488 | 0.0161 |
| 16 | T444 | M4-M4-M4 (crosslinked at D-Ala-DAP) | 2788.2020 | 2788.1819 | 0.0201 |
| 17 | D34$^{Anh}$ | M3-M4$^{Anh}$(crosslinked at DAP-DAP) | 1773.7415 | 1773.7339 | 0.0076 |
| 18 | D44$^{Anh}$ | M4-M4$^{Anh}$ (crosslinked at D-Ala-DAP) | 1844.7786 | 1844.7739 | 0.0047 |
| 19 | T444$^{Anh}$ | M4-M4-M4$^{Anh}$ (crosslinked at D-Ala-DAP) | 2768.1758 | 2768.1554 | 0.0204 |
| 20 | D44$^{2xAnh}$ | M4$^{Anh}$-M4$^{Anh}$ (crosslinked at D-Ala-DAP) | 1824.7524 | 1824.7451 | 0.0073 |

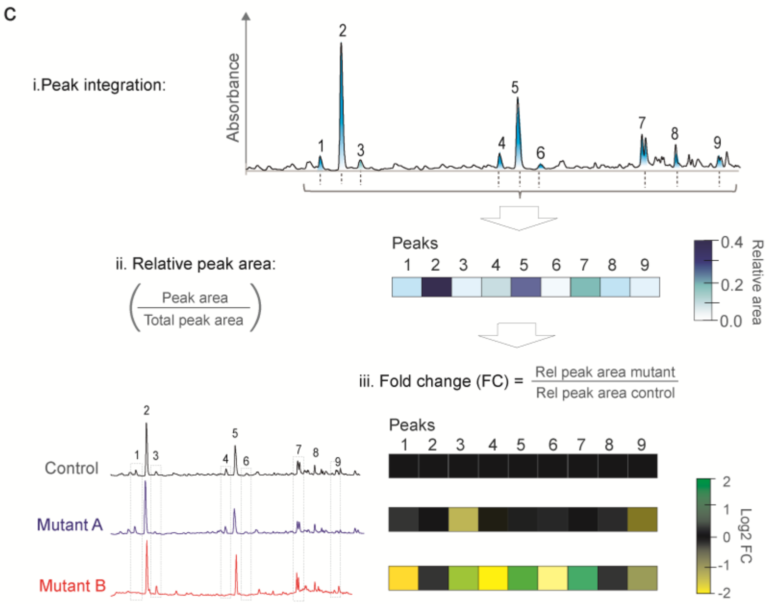


**Supplementary Fig. 2. Muropeptide profile of *V. cholerae*'s peptidoglycan obtained using the high throughput sample preparation method and analysis of data, Related to Figs. 2-3.**
**(a)** Representative chromatogram obtained for *V. cholerae* wildtype strain using the HT sample preparation method. Peaks of interest are listed. **(b)** Table of identified muropeptides in **a**. Identity was confirmed by MS/MS analysis, theoretical and observed neutral mass in Da are indicated. GlcNAc: N-acetyl glucosamine; MurNAc: N-acetyl muramic acid; (1-6anhydro) MurNAc: 1-6 anhydro N-acetyl muramic acid, terminal muropeptide; L-Ala: L-alanine; D-Glu: D-glutamic acid; DAP: *meso*-diaminopimelic acid; D-Ala: D-alanine; D-Met: D-methionine; Gly: glycine; Lpp: Braun's lipoprotein. **(c)** Pipeline for the data transformation and analysis: i) abundance was calculated by integration of the area of the peaks, ii) relative area of each muropeptide was calculated by dividing the peak area by the total area of the chromatogram, iii) finally, fold change (FC) relative to the control sample was calculated and Log2FC was used for representation in heat maps.

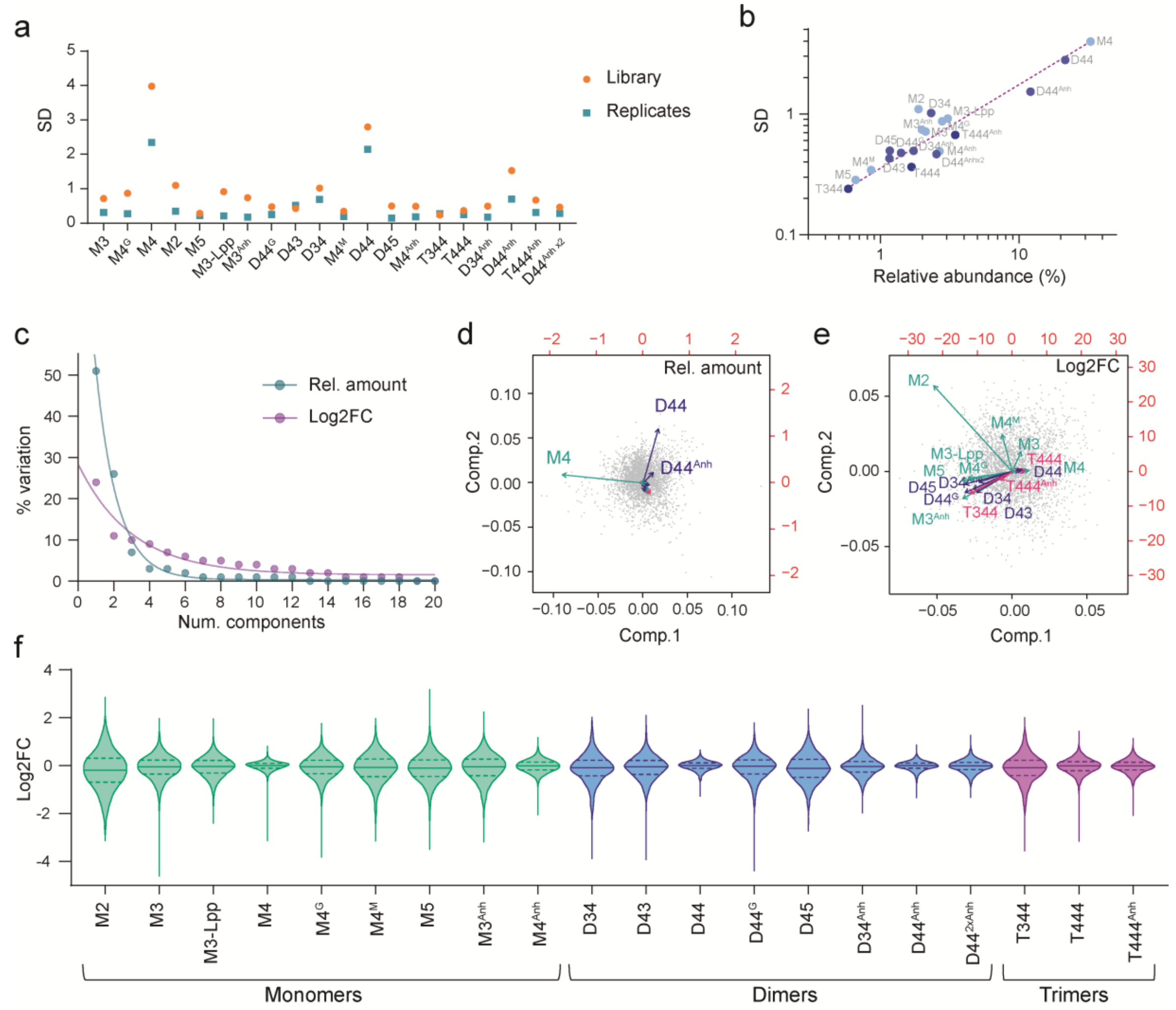


g

| Threshold | Mutants | % |
|---|---|---|
| 1SD | 2967 | 97.92 |
| 2SD | 1661 | 54.82 |
| 3SD | 581 | 19.17 |

h

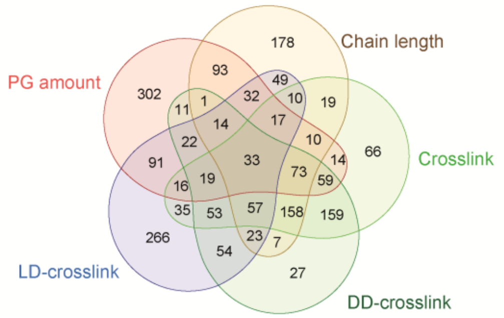


**Supplementary Fig. 3. Analysis of the peptidoglycan profiling of the *V. cholerae* non-redundant mutant library, Related to Fig. 3.**

(**a**) Biological versus technical variability: comparison of the standard deviation of the relative abundance of each muropeptide across all samples in the *V. cholerae* mutant library versus the average standard deviation across replicas of 300 samples. **(b)** Correlation between each muropeptide relative abundance and its standard deviation across the library dataset. **(c)** Scree plot of the main components used for the PCA using relative abundance of muropeptides and Log2FC calculated values. **(d)** Biplot showing the contribution of the different variables (muropeptides) to the data variability in the dataset of relative amounts**. (e)** Biplot showing the contribution of the different variables (muropeptides) to the data variability in the Log2FC dataset. **(f)** Violin plots showing the distribution of the samples for each muropeptide. Log2FC data was used. Median (continued lines) and quartiles (dashed lines) are represented. **(g)** Table representing the number and percentage of mutants with a significant phenotype for at least one muropeptide or a PG feature. A significant phenotype is considered when the sample is further than 1, 2 or 3 standard deviations (SD) from the average. **(h)** Venn diagram showing the number of outliers with a phenotype (using 1SD as threshold) in the main PG features.

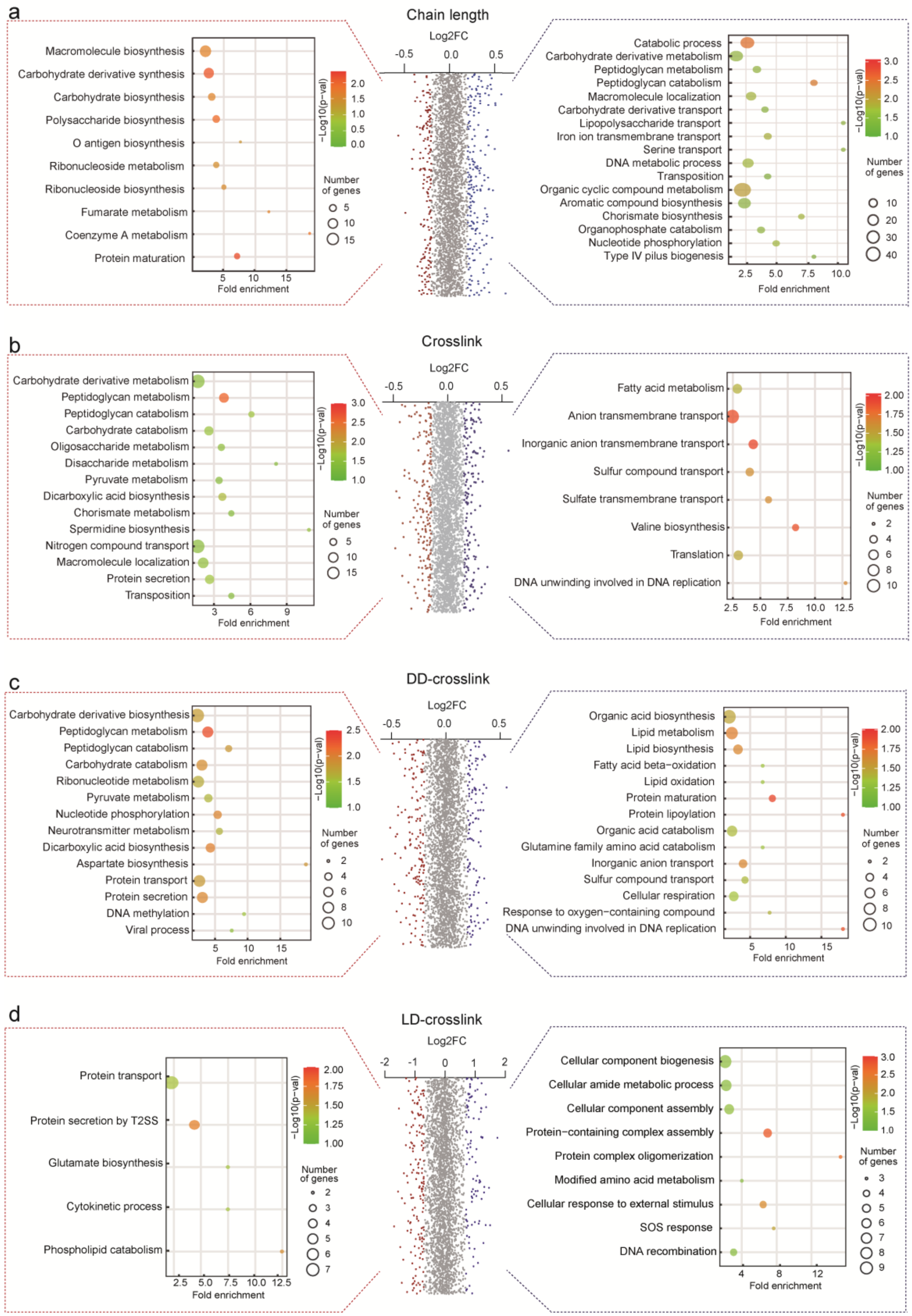


**Supplementary Fig. 4. Enrichment analysis of mutants presenting differential PG features, Related to Fig. 5.**
Over-represented biological processes of mutant candidates presenting low (left, red dots) and high (right, blue dots) relative amounts of: **(a)** chain length (Log2FC <-0.2 and >0.18); **(b)** total crosslink (Log2FC <-0.15 and >0.15); **(c)** DD-crosslink (Log2FC <-0.2 and >0.2); and **(d)** LD-crosslink (Log2FC <-0.7 and >0.7). The most representative biological processes with a fold enrichment higher than 2 are represented. The dot size in the balloon plots indicates the number of loci included in each category and the colour the significance of the enrichment (-Log10(p-values)).

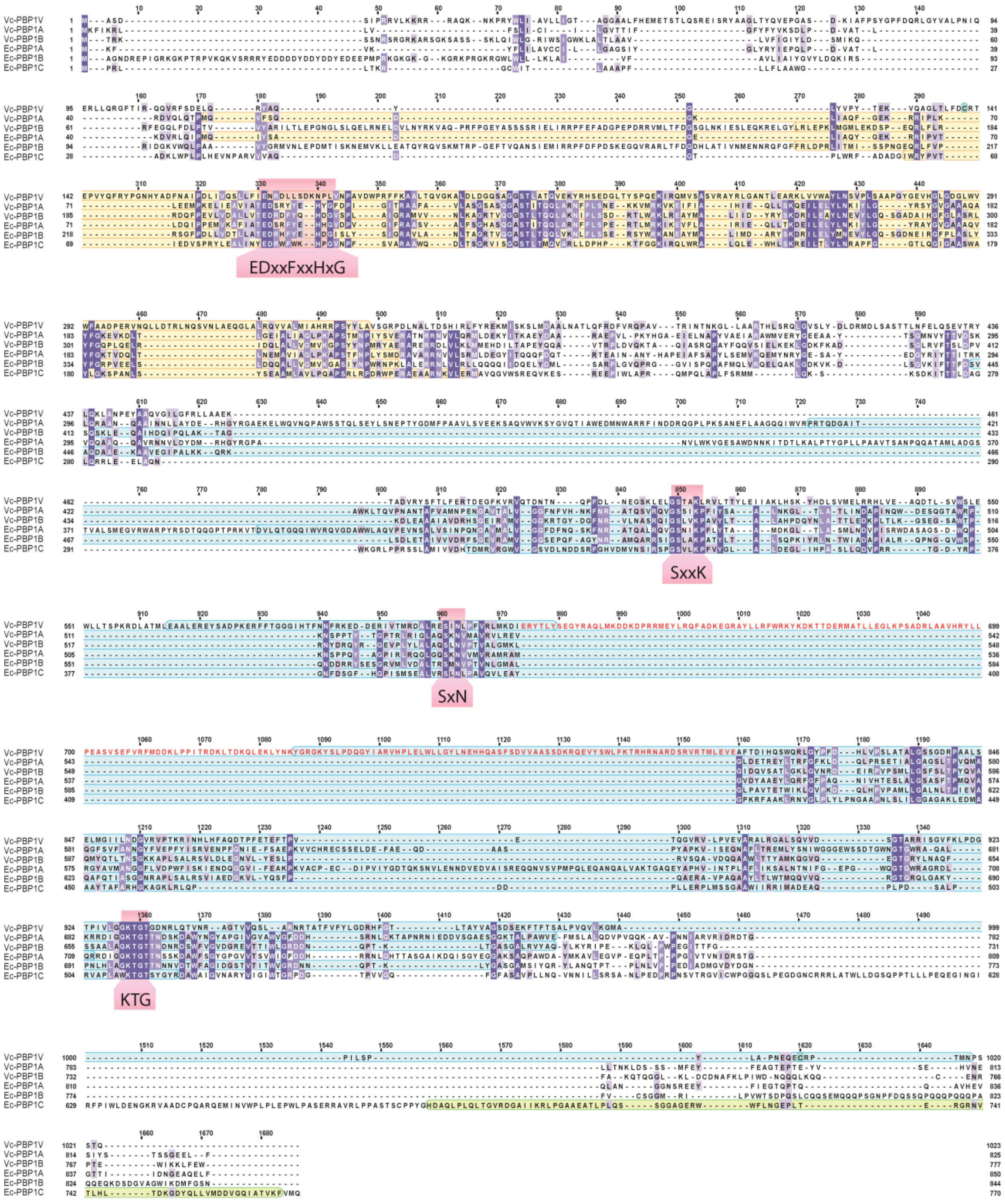


**Supplementary Fig. 5. Multiple sequence alignment of PBP1V with other high molecular weight penicillin-binding proteins, Related to Fig. 5.**

Sequence alignment of *V. cholerae* PBP1V (Vc-PBP1V, Uniprot: Q9KSD7), *V. cholerae* PBP1A (Vc-PBP1A, Uniprot: Q9KNU5), *V. cholerae* PBP1B (Vc-PBP1B, Uniprot: Q9KUC0), *E. coli* PBP1A (Ec-PBP1A, Uniprot: P02918), *E. coli* PBP1B (Ec-PBP1B, Uniprot: P02919) and *E. coli* PBP1C (Ec-PBP1C, Uniprot: A0A093EN65), performed with T-COFFEE Expresso. Background residue colour indicates degree of conservation. TG domain is highlighted in yellow, TP domain is highlighted in light blue, PBP1C-binding domain is highlighted in green. PBPs conserved motifs are indicated in the red boxes. Vc-PBP1V new domain in red letters.

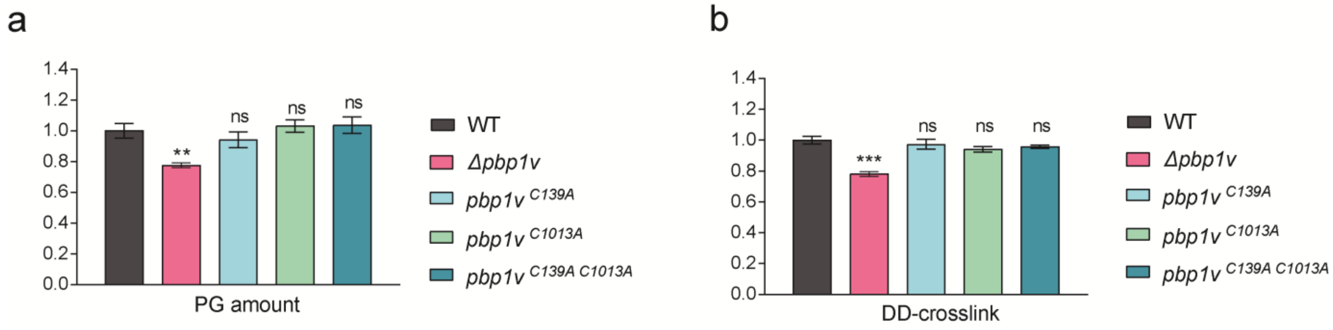


**Supplementary Fig. 6. Peptidoglycan analysis of PBP1V cysteine point mutants**, **Related to Fig. 5.** Quantification of the relative PG amount and DD-crosslink of the wildtype, Δ*pbp1V* mutant strain, and indicated cysteine mutants. (ns: not significant).
