## Extended Data 5 - Bacterial strains for "Genome-wide peptidoglycan profiling of *Vibrio cholerae*"

| Bacterial species | | Strain |  | |
| --- | --- | --- | --- | --- |
| *Vibrio cholerae* | | N16961 |  | |
| *Escherichia coli* | | K12 |  | |
| *Pseudomonas aeruginosa* | | PA14 |  | |
| *Salmonella enterica* serovar Typhimurium | | ATCC 14028 |  | |
| *Burkholderia thailandensis* | | PY79 |  | |
| *Acinetobacter baumannii* | | ATCC 17978 |  | |
| *Klebsiella pneumoniae* | | MKP103 |  | |
| *Bacillus subtilis* | | ATCC 6051 |  | |
| *Aeromonas hydrophila* | | ATCC 7966 |  | |
| *Burkholderia cenocepacia* | | K56-2 |  | |
| *Citrobacter rhodentium* | | ATCC 51116 |  | |
| *Enterobacter aerogenes* | | DSM 30053 |  | |
| *Enterobacter cloacae subsp. Cloacae* | | DSM 30054 |  | |
| *Photobacterium damselae* | | CIP102761 |  | |
| *Yersinia pseudotuberculosis* | | DSM 8992 |  | |
| *Enterococcus faecium* | | JH2-2 |  | |
| *Yersinia enterocolitica* | | DSM 4780 |  | |
| Bacteria | **Genotype** | | | **Source** |
| *V. cholerae* | *∆vc1321* | | | This study |
| *V. cholerae* | *vc1321 E172A* | | | This study |
| *V. cholerae* | *vc1321 S504A* | | | This study |
| *V. cholerae* | *vc1321 S610A* | | | This study |
| *V. cholerae* | *vc1321 C139A* | | | This study |
| *V. cholerae* | *vc1321 C1013A* | | | This study |
| *V. cholerae* | *vc1321 C1013A C139A* | | | This study |
| *V. cholerae* | *∆vc1321 pBAD18Km* | | | This study |
| *V. cholerae* | *∆vc1321 pBAD18Km::vc1321* | | | This study |
