## Extended Data 6 - Primers for "Genome-wide peptidoglycan profiling of *Vibrio cholerae*"

| Name | Sequence | Purpose |
| --- | --- | --- |
| vc1321-P1 | aaatctagagctttttacgtaaagtgacaattttg | Deletion mutant |
| vc1321-P2 | cttattgtgcttatatttattgcacttagttgaatccgtttaatcactg | Deletion mutant |
| vc1321-P3 | cggattcaactaagtgcaataaatataagcacaataag | Deletion mutant |
| vc1321-P4 | aaatctagacatatctaaaaaaacaatgagtgataacg | Deletion mutant |
| VC1321-pBADFOR | atcggaattcggagaggattcaactaagtggcttctg | pBAD overexpression |
| VC1321-pBADREV | tagctctagatgtgcttatatttattgtgtgc | pBAD overexpression |
| vc1321 E172A P1 | ccccctctagagttaaaaaagcgtcgacgcgc | Point mutant |
| vc1321 E172A P2 | ctcaacaaatcgcggttagcgatgaacaatagcgac | Point mutant |
| vc1321 E172A P3 | gtcgctattgttcatcgctaaccgcgatttgttgag | Point mutant |
| vc1321 E172A P4 | ccccctctagacgacccgatacggctagg | Point mutant |
| vc1321 S504A P1 | aaaatctagagtgtgggcttatttgaac | Point mutant |
| vc1321 S504A P2 | tacacgcagcttagcggtagcgcccaactccaactt | Point mutant |
| vc1321 S504A P3 | aagttggagttgggcgctaccgctaagctgcgtgta | Point mutant |
| vc1321 S504A P4 | aaaatctagaccgagcgatataaccttg | Point mutant |
| vc1321 S610A P1 | ccccctctagatcttggtttccgtttactggc | Point mutant |
| vc1321 S610A P2 | aaaggcaggttgatagcctcacgcaatgcatca | Point mutant |
| vc1321 S610A P3 | tgatgcattgcgtgaggctatcaacctgccttt | Point mutant |
| vc1321 S610A P4 | ccccctctagagcgatataaccttgatccggc | Point mutant |
| vc1321 C139A P1 | aaaatctagacttctacactattatcatcgg | Point mutant |
| vc1321 C139A P2 | taaacgggttcagtacgagcatcaaataaggtaag | Point mutant |
| vc1321 C139A P3 | cttaccttatttgatgctcgtactgaacccgttta | Point mutant |
| vc1321 C139A P4 | aaaatctagactgattgagtcgagtatcc | Point mutant |
| vc1321 C1013A P1 | aaaatctagaggtagctcgggtgatag | Point mutant |
| vc1321 C1013A P2 | ggttcatcgtaggtcgagcctcttgctcattcgg | Point mutant |
| vc1321 C1013A P3 | ccgaatgagcaagaggctcgacctacgatgaacc | Point mutant |
| vc1321 C1013A P4 | aaaatctagacacagttacagccaatcac | Point mutant |
